# *GWAS-Flow*: A GPU accelerated framework for efficient permutation based genome-wide association studies

**DOI:** 10.1101/783100

**Authors:** Jan A. Freudenthal, Markus J. Ankenbrand, Dominik G. Grimm, Arthur Korte

## Abstract

**Motivation:** Genome-wide association studies (GWAS) are one of the most commonly used methods to detect associations between complex traits and genomic polymorphisms. As both genotyping and phenotyping of large populations has become easier, typical modern GWAS have to cope with massive amounts of data. Thus, the computational demand for these analyses grew remarkably during the last decades. This is especially true, if one wants to implement permutation-based significance thresholds, instead of using the naïve Bonferroni threshold. Permutation-based methods have the advantage to provide an adjusted multiple hypothesis correction threshold that takes the underlying phenotypic distribution into account and will thus remove the need to find the correct transformation for non Gaussian phenotypes. To enable efficient analyses of large datasets and the possibility to compute permutation-based significance thresholds, we used the machine learning framework TensorFlow to develop a linear mixed model (`GWAS-Flow`) that can make use of the available CPU or GPU infrastructure to decrease the time of the analyses especially for large datasets.

**Results:** We were able to show that our application `GWAS-Flow` outperforms custom GWAS scripts in terms of speed without loosing accuracy. Apart from p-values, `GWAS-Flow` also computes summary statistics, such as the effect size and its standard error for each individual marker. The CPU-based version is the default choice for small data, while the GPU-based version of `GWAS-Flow` is especially suited for the analyses of big data.

**Availability:** `GWAS-Flow` is freely available on GitHub (https://github.com/Joyvalley/GWAS_Flow) and is released under the terms of the MIT-License.

## Introduction

Genome-wide association studies, pioneered in human genetics [1] in the last decade, have become the predominant method to detect associations between phenotypes and the genetic variations present in a population. Understanding the genetic architecture of traits and mapping the underlying genomic polymorphisms is of paramount importance for successful breeding both in plants and animals, as well as for studying the genetic risk factors of diseases. Over the last decades, the cost for genotyping have been reduced dramatically. Early GWAS consisted of a few hundred individuals which have been phenotyped and genotyped on a couple of hundreds to thousands of genomic markers. Nowadays, marker density for many species easily exceed millions of genomic polymorphisms. Albeit commonly SNPs are used for association studies, standard GWAS models are flexible to handle different genomic features as input. The *Arabidopsis* 1001 genomes project features for example 1135 sequenced *Arabidopsis thaliana* accessions with over 10 million genomic markers that segregate in the population [2]. Other genome projects also yielded large amounts of genomic data for a substantial amount of individuals, as exemplified in the 1000 genomes project for humans [3], the 2000 yeast genomes project or the 3000 rice genomes project [4]. Thus, there is an increasing demand for GWAS models that can analyze these data in a reasonable time frame. One critical step of GWAS is to determine the threshold at which an association is termed significant. Classically the conservative Bonferroni threshold is used, which accounts for the number of statistical tests that are performed, while many recent studies try to use significance thresholds that are based on the false-discovery rate (FDR) [5]. An alternative approach are permutation-based thresholds [6]. Permutation-based thresholds estimate the significance by shuffling phenotypes and genotypes before each GWAS run, thus any signal left in the data should not have a genetic cause, but might represent model mis-specifications or uneven phenotypic distributions. Typically this process is repeated hundreds to thousands of times and will lead to a distinct threshold for each phenotype analyzed [7]. The computational demand of permutation-based thresholds is immense, as per analysis not one, but at least hundreds of GWAS need to be performed. Here the main limitation is the pure computational demand. Thus, faster GWAS models could easily make the estimation of permutation-based thresholds the default choice.

## Methods

### GWAS Model

The GWAS model used for `GWAS-Flow` is based on a fast approximation of the linear-mixed-model described in [8, 9], which estimates the variance components *σ*_g_ and *σ*_e_ only once in a null model that includes the genetic relationship matrix, but no distinct genetic markers. These components are thereafter used for the tests of each specific marker. Here, the underlying assumption is, that the ratio of these components stays constant, even if distinct genetic markers are included into the GWAS model. This holds true for nearly all markers and only markers which posses a big effect will alter this ratio slightly, where now *σ*_g_ would become smaller compared to the null model. Thus, the p-values calculated by the approximation might be a little higher (less significant) for strongly associated markers.

### The GWAS-Flow Software

The `GWAS-Flow` software was designed to provide a fast and robust GWAS implementation that can easily handle large data and allows to perform permutations in a reasonable time frame. Traditional GWAS implementations that are implemented using Python [10] or R [11] cannot always meet these demands. We tried to overcome those limitations by using TensorFlow [12], a multi-language machine learning framework published and developed by Google. GWAS calculations are composed of a series of matrix computations that can be highly parallelized, and easily integrated into the architecture provided by TensorFlow. Our implementation allows both, the classical parallelization of code on multiple processors (CPUs) and the use of graphical processing units (GPUs). `GWAS-Flow` is written using the Python TensorFlow API. Data import is done with *pandas* [13] *and/or HDF5* for Python [14]. Preprocessing of the data (e.g filtering by minor Allele count (MAC)) is performed with *numpy* [15]. Variance components for residual and genomic effects are estimated with a slightly altered function based on the Python package *limix* [16]. The GWAS model is based on the following linear mixed model that takes into account the effect of every marker with respect to the kinship:

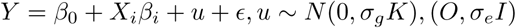

The residuals are used to calculate a p-value for each marker according to an overall F-test that compares the model including a distinct genetic effect to a model without this genetic effect:

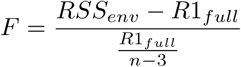

Apart from the p-values that derive from the F-distribution, `GWAS-Flow` also report summary statistics, such as the estimated effect size (*β*_*i*_) and its standard error for each marker.

### Calculation of the permutation-based threshold for GWAS

To calculate a permuation-based threshold, we essentially perform *n* repetitions (*n* > 100) of the GWAS on the same data with the sole difference that before each GWAS we randomize the phenotypic values. Thus any correlation between the phenotype and the genotype will be broken and indeed for over 90% of these analyses the estimated pseudo-heritability is close to zero. On the other hand, the phenotypic distribution will stay unaltered by this randomization. Hence, any remaining signal in the GWAS has to be of a non-genetic origin and could be caused by e.g. model mis-specifications. Now we take the lowest p-value (after filtering for the desired minor allele count) for each permutation and take the 5% lowest value as the permutation-based threshold for the GWAS.

### Dissemination and reproducibility

`GWAS-Flow` is an open-source software and was published on GitHub (https://github.com/Joyvalley/GWAS_Flow) under the terms of the MIT-License making `GWAS-Flow` free to use and alter for the scientific community. All calculations mentioned in the study were performed with the first stable version v1.0. Detailed installation information are given in the `README.md` file on GitHub. We provide three different ways to install and run `GWAS-Flow`: (i) with virtual environments using Anaconda for Python, (ii) Docker containers that in version 1.0 have the exact setup used for the calculations in this study to ensure full reproducibility [17]. Besides Docker no other Software is required. (iii) To make use of the advantages of containerized solutions in multi-user HPC environments we also provide instructions for compilation of singularity images [18].

### Benchmarking

For benchmarking of `GWAS-Flow` we used data from the *Arabidopsis* 1001 Genomes Project [2]. The genomic data we used were subsets between 10,000 and 100,000 markers. We chose not to include subsets that exceed 100,000 markers, because there is a linear relationship between the number of markers and the computational time demanded, as all markers are tested independently. We used phenotypic data for flowering time at ten degrees (FT10) for *A. thaliana*, published and downloaded from the AraPheno database [19]. We down- and up-sampled sets to generate phenotypes for sets between 100 and 5000 accessions. For each set of phenotypes and markers we ran 10 permutations to assess the computational time needed. All analyses have been performed with a custom R script that has been used previously [7], `GWAS-Flow` using either a CPU or a GPU architecture and *GEMMA* [20]. *GEMMA* is a fast and efficient implementation of the mixed model that is broadly used to perform GWAS. All calculations were run on the same machine using 16 i9 virtual CPUs. The GPU version ran on an NVIDIA Tesla P100 graphic card. Additionally to the analyses of the simulated data, we compared the times required by *GEMMA* and both `GWAS-Flow` implementations for > 200 different real datasets from *A. thaliana* that have been downloaded from the AraPheno [19] database and have been analyzed with the available fully imputed genomic dataset of ∼10 million markers, filtered for a minor allele count greater five.

## Results

The two main factors influencing the computational time for GWAS are the number of markers incorporated in such an analysis and the number of different accessions, while the latter has an approximate quadratic effect in classical GWAS implementations [20]. Figure 1A shows the time demand as a function of the number of accessions used in the analysis with 10,000 markers. The quadratic increase in time demand is clearly visible for the custom R implementation, as well as for the CPU-based `GWAS-Flow` implementation and *GEMMA*. The `GWAS-Flow` implementation and *GEMMA* clearly outperforms the R implementation in general, while for a small number of accessions `GWAS-Flow` is slightly faster then *GEMMA*. For the GPU-based implementation the increase in run-time with larger sample sizes is much less pronounced. While for small (< 1,000 individuals) data, there is no benefit compared to running `GWAS-Flow` on CPUs or running *GEMMA*, the GPU-version clearly outperforms the other implementations if the number of accessions increases. Figure 1B shows the computational time in relation to the number of markers and a fixed amount of 2000 accessions for the two different `GWAS-Flow` implementations. Here, a linear relationship is visible in both cases. To show the performance of `GWAS-Flow` not only for simulated data, we also run both implementations on more than 200 different real datasets downloaded from the AraPheno database. Figure 1C shows the computational time demands for all analyses comparing both `GWAS-Flow` implementation to *GEMMA*. Here, the CPU-based `GWAS-Flow` performs comparable to *GEMMA*, while the GPU-based implementation outperforms both, if the number of accessions is above 500. Importantly all obtained GWAS results (p-values, beta estimates and standard errors of the beta estimates) are nearly (apart from some mathematical inaccuracies) identical between the three different implementations.

**Figure 1.**
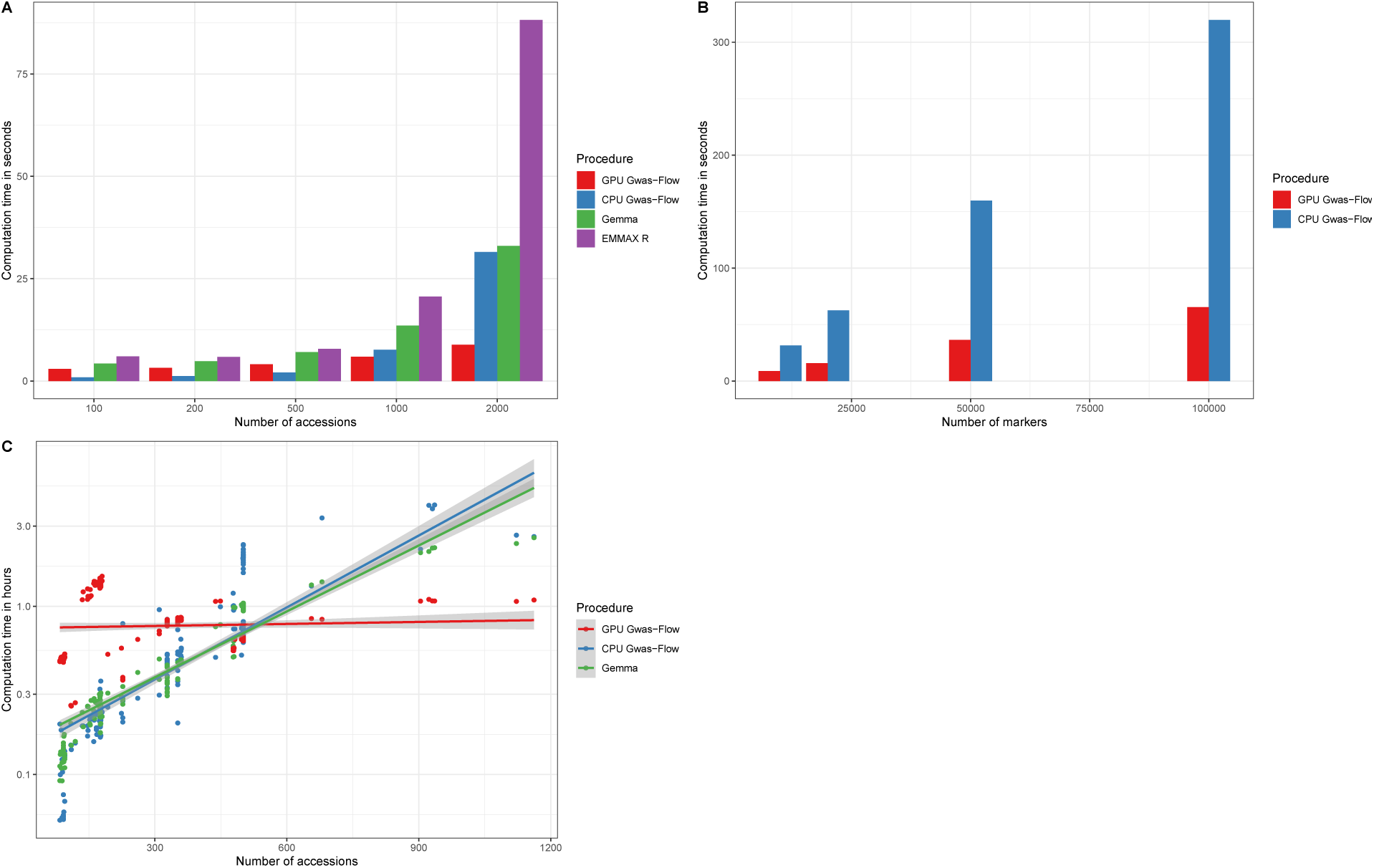
Comparison of the computational time needed for GWAS between GWAS-Flow, *GEMMA* and a custom R script. **A** Computational time as a function of the number of accessions with 10000 markers each. **B** Computational time as a function of the number of genetic markers with constantly 2000 accessions for both `GWAS-Flow` versions. **C** Comparison of the computational time for the analyses of > 200 phenotypes from *Arabidopsis thaliana* as a function of the number of accessions for *GEMMA* and the CPU- and GPU-based version of `GWAS-Flow`. GWAS was performed with a fully imputed genotype matrix containing 10.7 M markers and a minor allele filter of MAC > 5.

## Discussion

We made use of recent developments of computational architecture and software to cope with the increasing computational demand in analyzing large GWAS datasets. With `GWAS-Flow` we implemented both, a CPU- and a GPU-based version of the classical linear mixed model commonly used for GWAS. Both implementations outperform custom R scripts on simulated and real data. While the CPU-based version performs nearly identical compared to *GEMMA*, a commonly used GWAS implementation, the GPU-based implementation outperforms both, if the number of individuals, which have been phenotyped, increases. For analyzing big data, here the main limitation would be the RAM of the GPU, but as the individual test for each marker are independent, this can be easily overcome programmatically. The presented `GWAS-Flow` implementations are markedly faster compared to custom GWAS scripts and even outperform efficient fast implementations like *GEMMA* in terms of speed. This readily enables the use of permutation-based thresholds, as with `GWAS-Flow` hundred permutations can be performed in a reasonable time even for big data. Thus, it is possible for each analyzed phenotype to create a specific, permutation-based threshold that might present a more realistic scenario. Importantly the permutation-based threshold can be easily adjusted to different minor allele counts, generating different significance thresholds depending on the allele count. This could help to distinguish false and true associations even for rare alleles. `GWAS-Flow` is a versatile and fast software package. Currently `GWAS-Flow` is and will remain under active development to make the software more versatile. This will e.g. include the compatibility with TensorFlow v2.0.0 and enable data input formats, such as PLINK [21]. The whole framework is flexible, so it is easy to include predefined co-factors e.g to enable multi-locus models [22] or account for multi-variate models like the multi-trait mixed model [23]. Standard GWAS are good in detecting additive effects with comparably large effect sizes, but lack the ability to detect epistatic interactions and their influence on complex traits [24, 25]. To catch the effects of these gene-by-gene or SNP-by-SNP interactions, a variety of genome-wide association interaction studies (GWAIS) have been developed, thoroughly reviewed in [26]. Here, `GWAS-Flow` might provide a tool that enables to test the full pairwise interaction matrix of all SNPs. Although this might be a statistic nightmare, it now would be computationally feasible.

## Funding

Funding from the BMBF to A.K.(grant 031B0195H).

## Acknowledgments

We thank Pamela Korte for proof-reading of the manuscript.

## Notes

https://github.com/Joyvalley/GWAS_Flow

## References

1. J. N. Hirschhorn and M. J. Daly, “Genome-wide association studies for common diseases and complex traits,” Nature Reviews Genetics, vol. 6, no. 2, pp. 95–108, 2005.

2. C. Alonso-Blanco, J. Andrade, C. Becker, F. Bemm, J. Bergelson, K. M. Borgwardt, J. Cao, E. Chae, T. M. Dezwaan, W. Ding, et al., “1,135 genomes reveal the global pattern of polymorphism in arabidopsis thaliana,” Cell, vol. 166, no. 2, pp. 481–491, 2016.

3. N. Siva, “1000 genomes project,” 2008.

4. J.-Y. Li, J. Wang, and R. S. Zeigler, “The 3,000 rice genomes project: new opportunities and challenges for future rice research,” GigaScience, vol. 3, no. 1, p. 8, 2014.

5. J. D. Storey and R. Tibshirani, “Statistical significance for genomewide studies,” Proceedings of the National Academy of Sciences, vol. 100, no. 16, pp. 9440–9445, 2003.

6. R. Che, J. R. Jack, A. A. Motsinger-Reif, and C. C. Brown, “An adaptive permutation approach for genome-wide association study: evaluation and recommendations for use,” BioData mining, vol. 7, no. 1, p. 9, 2014.

7. M. Togninalli, Ü. Seren, D. Meng, J. Fitz, M. Nordborg, D. Weigel, K. Borgwardt, A. Korte, and D. G. Grimm, “The aragwas catalog: a curated and standardized arabidopsis thaliana gwas catalog,” Nucleic acids research, vol. 46, no. D1, pp. D1150–D1156, 2017.

8. H. M. Kang, J. H. Sul, S. K. Service, N. A. Zaitlen, S.-y. Kong, N. B. Freimer, C. Sabatti, E. Eskin, et al., “Variance component model to account for sample structure in genome-wide association studies,” Nature genetics, vol. 42, no. 4, p. 348, 2010.

9. Z. Zhang, E. Ersoz, C.-Q. Lai, R. J. Todhunter, H. K. Tiwari, M. A. Gore, P. J. Bradbury, J. Yu, D. K. Arnett, J. M. Ordovas, and E. S. Buckler, “Mixed linear model approach adapted for genome-wide association studies,” Nature Genetics, vol. 42, pp. 355–360, Mar. 2010.

10. G. Van Rossum and F. L. Drake Jr, Python tutorial. Centrum voor Wiskunde en Informatica Amsterdam, The Netherlands, 1995.

11. R Core Team, R: A Language and Environment for Statistical Computing. R Foundation for Statistical Computing, Vienna, Austria, 2019.

12. M. Abadi, A. Agarwal, P. Barham, E. Brevdo, Z. Chen, C. Citro, G. S. Corrado, A. Davis, J. Dean, M. Devin, S. Ghemawat, I. Goodfellow, A. Harp, G. Irving, M. Isard, Y. Jia, R. Jozefowicz, L. Kaiser, M. Kudlur, J. Levenberg, D. Mané, R. Monga, S. Moore, D. Murray, C. Olah, M. Schuster, J. Shlens, B. Steiner, I. Sutskever, K. Talwar, P. Tucker, V. Vanhoucke, V. Vasudevan, F. Viégas, O. Vinyals, P. Warden, M. Wattenberg, M. Wicke, Y. Yu, and X. Zheng, “TensorFlow: Large-scale machine learning on heterogeneous systems,” 2015. Software available from tensorflow.org.

13. W. McKinney, “Data structures for statistical computing in python,” in Proceedings of the 9th Python in Science Conference (S. van der Walt and J. Millman, eds.), pp. 51 – 56, 2010.

14. A. Collette, Python and HDF5. O’Reilly, 2013.

15. T. E. Oliphant, A guide to NumPy, vol. 1. Trelgol Publishing USA, 2006.

16. C. Lippert, F. P. Casale, B. Rakitsch, and O. Stegle, “Limix: genetic analysis of multiple traits,” bioRxiv, 2014.

17. D. Merkel, “Docker: lightweight linux containers for consistent development and deployment,” Linux Journal, vol. 2014, no. 239, p. 2, 2014.

18. G. M. Kurtzer, V. Sochat, and M. W. Bauer, “Singularity: Scientific containers for mobility of compute,” PloS one, vol. 12, no. 5, p. e0177459, 2017.

19. Ü. Seren, D. Grimm, J. Fitz, D. Weigel, M. Nordborg, K. Borgwardt, and A. Korte, “Arapheno: a public database for arabidopsis thaliana phenotypes,” Nucleic acids research, p. gkw986, 2016.

20. X. Zhou and M. Stephens, “Genome-wide efficient mixed-model analysis for association studies,” Nature Genetics, vol. 44, pp. 821–824, June 2012.

21. S. Purcell, B. Neale, K. Todd-Brown, L. Thomas, M. A. Ferreira, D. Bender, J. Maller, P. Sklar, P. I. De Bakker, M. J. Daly, et al., “Plink: a tool set for whole-genome association and population-based linkage analyses,” The American journal of human genetics, vol. 81, no. 3, pp. 559–575, 2007.

22. V. Segura, B. J. Vilhjálmsson, A. Platt, A. Korte, Ü. Seren, Q. Long, and M. Nordborg, “An efficient multi-locus mixed-model approach for genome-wide association studies in structured populations,” Nature genetics, vol. 44, no. 7, p. 825, 2012.

23. A. Korte, B. J. Vilhjálmsson, V. Segura, A. Platt, Q. Long, and M. Nordborg, “A mixed-model approach for genome-wide association studies of correlated traits in structured populations,” Nature genetics, vol. 44, no. 9, p. 1066, 2012.

24. B. Mckinney and N. Pajewski, “Six degrees of epistasis: statistical network models for gwas,” Frontiers in genetics, vol. 2, p. 109, 2012.

25. A. Korte and A. Farlow, “The advantages and limitations of trait analysis with gwas: a review,” Plant methods, vol. 9, no. 1, p. 29, 2013.

26. M. D. Ritchie and K. Van Steen, “The search for gene-gene interactions in genome-wide association studies: challenges in abundance of methods, practical considerations, and biological interpretation,” Annals of translational medicine, vol. 6, no. 8, 2018.

